# Strategies for Mitigating Radiation Damage and Improving Data Completeness in 3D Electron Diffraction of Protein Crystals

**DOI:** 10.1101/2025.02.06.636927

**Authors:** Alaa Shaikhqasem, Farzad Hamdi, Lisa Machner, Christoph Parthier, Constanze Breithaupt, Fotis L. Kyrilis, Stephan M. Feller, Panagiotis L. Kastritis, Milton T. Stubbs

**Author notes:** correspondence to: Panagiotis L. Kastritis or Milton T. Stubbs. These authors contributed equally to this work (Alaa Shaikhqasem, Farzad Hamdi).

## Abstract

While 3D electron diffraction (3D-ED or microcrystal electron diffraction, MicroED) has emerged as a promising method for protein structure determination, its applicability is hindered by high susceptibility to radiation damage, leading to decreasing signal-to-noise ratio in consecutive diffraction patterns that limits the quality (resolution and redundancy) of the data. In addition, data completeness may be restricted due to the geometrical limitations of current sample holders and stages. In this work, we introduce an approach that addresses these issues using a commonly available 200keV cryo-electron microscope. The multi-position acquisition technique we present here combines (a) multiple data acquisitions from a single crystal over several tilt ranges and (b) merging data from a small number of crystals, each tilted about a different axis. The robustness of this approach is demonstrated by the *de novo* elucidation of a protein-peptide complex structure from only two orthorhombic microcrystals.

## Introduction

X-ray crystallography has represented a major workhorse for biological macromolecular structure determination for over six decades. The low X-ray scattering cross-section of light atoms typically requires the use of macroscopic (**≳** 10^−4^ m) crystals to achieve sufficient diffracting volumes. Initial macromolecule crystallization screening however often results in microcrystals, needle-like structures, needle clusters and/or inhomogeneous crystals with limited X-ray diffracting power. While extensive fine screening of buffer conditions may be used to obtain large single crystals, this is by no means a certain outcome. The development of synchrotron microfocus beamlines has allowed the collection of X-ray diffraction data suitable for structure determination from micro- and imperfect crystals [1]. Recent technological advances in X-ray crystallography (notably the development of serial femtosecond nanocrystallography (SFX) and the emergence of X-ray free electron lasers (XFELs)) have opened new possibilities for collecting X-ray diffraction data from sub-micron protein crystals [2]–[5]. For these techniques, a continuous flow of thousands of microcrystals is directed into the path of the X-ray beam and individual diffraction frames are collected from randomly oriented crystals [6], [7]. Nonetheless, the production of a jet of microcrystals is technically challenging, frames from several thousand sub-micron crystals must be collected, scaled and merged together to generate a complete dataset [8], [9], and widespread adoption of these techniques is hindered by limited access to relevant facilities [2].

As electrons interact with matter far more strongly than X-rays, in house electron diffraction (ED) represents a promising alternative. Indeed, ED of unstained sub-micron two dimensional crystals was used to elucidate the first three dimensional model of a membrane protein (bacteriorhodopsin) already half a century ago [10]. The short electron wavelength (∼pm = 10^−2^ Å, which results in a flat Ewald sphere) complicates the indexing of ED patterns from three dimensional crystals, however [11]. This has been mitigated through the development of continuous rotation diffraction data collection techniques, termed three-dimensional electron diffraction (3D-ED) or microcrystal electron diffraction (MicroED) [12], [13]. The high electron scattering cross section can however lead to (i) substantial inelastic scattering, which lowers the signal-to-noise ratio of the diffracted data and (ii) multiple scattering events (“dynamical diffraction”) from successive layers of the crystal, which complicates the relationship between the Coulomb potential density and the acquired intensities. To reduce both effects, thin (⪅ 400 nm) crystals are preferred for ED analyses [14].

Substantial progress in 3D-ED/MicroED over the past decade [15] has allowed the determination of protein structures from crystals of micrometer and sub-micrometer dimensions [13], [16]– [18], with more than a hundred protein structures now deposited in the Protein Data Bank (PDB, https://www.rcsb.org/stats). Most of these depositions however correspond to model proteins that have been studied to explore the potential and limitations of 3D-ED/MicroED as well as for methodological developments [19]–[32]. Only in 2019 was the first novel protein structure – that of the metalloenzyme R2lox – determined by 3D-ED/MicroED [33], which was achieved by merging data collected from 21 crystals to 3 Å resolution with a completeness (the ratio between the number of measured and theoretically observable reflections) of 62.8 %.

In a 3D-ED/MicroED analysis of biological macromolecules, small micro- or nanometer-sized crystals (microcrystals) are prepared by applying and vitrifying the sample on an EM grid, from which diffraction data are collected while rotating the sample stage continuously under a low-dose rate electron beam [18]. Recorded diffraction images are processed using conventional X-ray crystallography data processing software, and Fourier transformation of the phased data set yields a Coulomb potential map that can be used for model building [34], [35]. A major limitation of the method is the accumulation of radiation damage over the course of data acquisition [19], [29], which has an adverse effect on data quality, resolution and completeness. Reduced data completeness, which can greatly impact map quality and model accuracy [36], also results from the limited rotation range of transmission electron microscope (TEM) stages, and can be further affected by crystal location on the grid, sample quality and crystal-specific characteristics such as shape and symmetry. While merging data from multiple randomly oriented crystals can be employed to enhance data completeness [22], [29], [37], [38], this demands a high number of crystals. Serial electron crystallography (Serial-ED), whereby single ED patterns are collected from randomly oriented crystals on a sample grid at a fixed tilt angle, can address some of these issues [39], [22], although (i) it is in general not possible to index protein diffraction data from a single frame due to the flatness of the Ewald sphere, necessitating prior knowledge of the space groups for data processing, and (ii) processing a complete 3D-ED/MicroED data set is dependent on merging data from thousands of crystals. The latter is particularly problematic in cases where the crystals are heterogeneous in their packing (“non-isomorphism”).

Thus, there is a need for a better data acquisition strategy that can be applied to samples with a small number of diffracting crystals. This paper presents such an optimized data acquisition and processing strategy for the collection of high-quality electron diffraction data from extended crystals using conventional cryo-electron microscopy (cryo-EM) instrumentation. Our approach, inspired by helical data acquisition schemes employed in X-ray crystallography [40], [41], yields both high data completeness and redundancy. This is achieved by collecting and merging angular segments of ED data from non-contiguous regions of a single crystal, which are then merged with corresponding data from other differently oriented crystals. This multi-position acquisition strategy provides an effective means of reducing any effects of non-isomorphism while minimizing radiation damage, allowing acquisition of complete electron diffraction data from beam-sensitive crystals. We have applied this method to solve the structure of the C-terminal peptide (amino acids 617-684) of human Grb2-associated binding protein 1 (Gab1^617-684^) in complex with the N-terminal region (amino acids 1-222) of the tyrosine-protein phosphatase non-receptor type 11 (SHP2^1-222^). We also discuss here the challenges posed by sample preparation and data processing for structure determination as well as our solutions, which can be further fine-tuned on a case-by-case basis for other samples.

## Materials and Methods

### Preliminary experiments

Initial electron diffraction experiments utilizing our in-house 200 kV instrument were performed following published protocols for lysozyme [13], [22], [25], [29], [42] (Supplementary Methods S1). The imaging protocols applied are given in Supplementary Methods S2.

### Sample Preparation

Protein expression and purification of SHP2^1-222^ and the intrinsically disordered C-terminal part of Gab1^617-684^ phosphorylated at Tyr 627 and Tyr 659 are described elsewhere [43]. Prior to crystallization, SHP2^1-222^ (25 mg/ml) was mixed with Gab1^617-684^ (7.7 mg/ml) in 20 mM Bis-Tris pH 6.5, 50 mM NaCl, 1 mM DTT. The complex was crystallized by hanging drop vapor diffusion in 15 well EasyXtal crystallization plates (Qiagen). Thin needle-shaped crystals appeared in 0.1 M Tris pH 8.8, 29% PEG3350 within seven days at 12 °C (Figure 1A), which were followed after another ten days by more compact plate-like crystals (Figure 1B).

**Figure 1:**
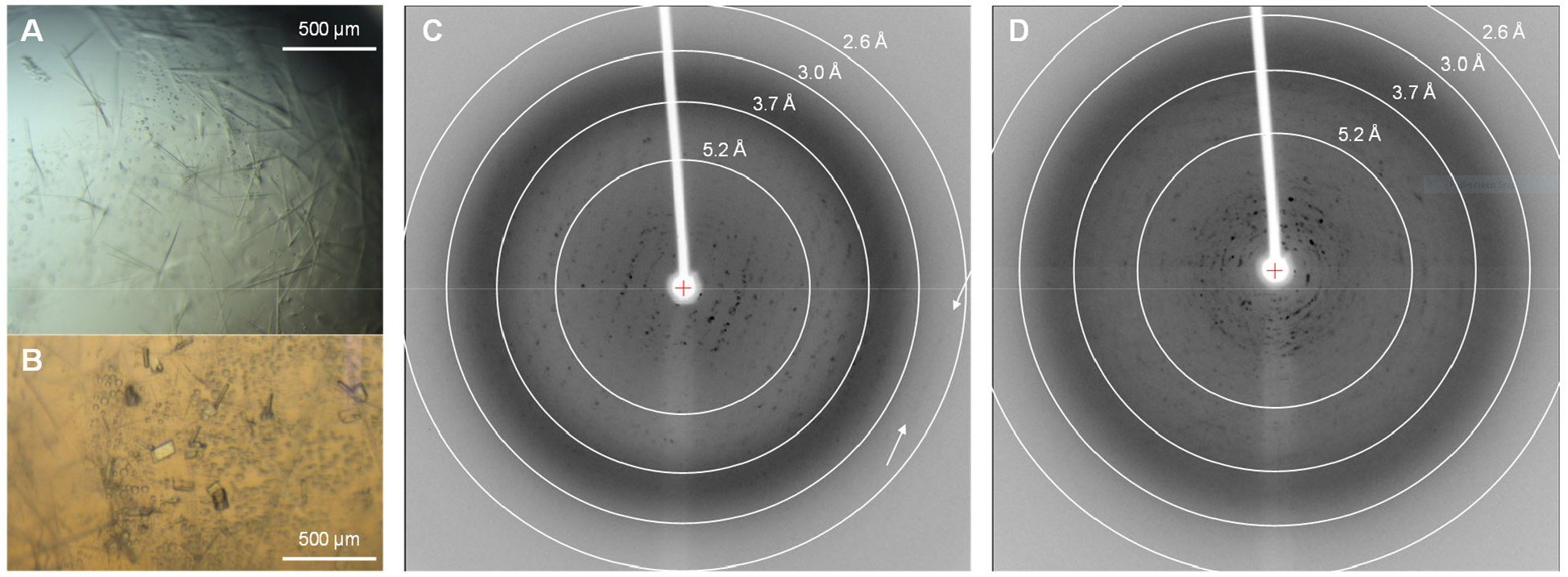
Macroscopic crystals of the Gab1-SHP2 complex demonstrate disorder. (A) Light microscope view of needle-like Gab1-SHP2 crystals grown in a hanging drop using the vapor diffusion method. (B) After 18 days, small plate-like crystals grew in the same drop. (C, D) X-ray diffraction patterns of two independent plate-like crystals (B) reveal Bragg reflections up to around 2.7 Å (white arrows), but exhibit multiple lattices and high mosaicity.

For 3D-ED/MicroED, a skin that formed at the surface of the drop was removed and the remaining drop, including the mother liquor and the crystals, was transferred to a 1.5 ml tube. Crystals were pooled from several crystallization drops and used directly to make vitrified samples. Quantifoil® R2/1 grids, plasma treated in a PELCO easiGLOW glow discharging machine at 15 mA for 25 seconds under 0.4 mBar residual air pressure, were mounted in a ThermoFisher Scientific Vitrobot Mark IV (which is designed to blot both sides simultaneously (front and back)) with chamber temperature and humidity adjusted to 4 °C and > 95 % respectively. Sample vitrification was modified from that used for lysozyme as outlined in Supplementary Methods S3. The filter paper facing the sample (front face) was replaced by a similarly cut disc of clean and unstretched parafilm (see Supplementary Figure S3E) and a 3.5 µl aliquot of the crystal-containing suspension applied to the carbon film side of the grid (front face). A 3.5 µl of reservoir solution mixed with water (1:1 ratio) was applied to the other (back) face of the grid facing a 595-grade ashless filter paper pad and the grid blotted for 25 seconds, causing the excess liquid to be absorbed from the back face and the crystals to settle on the carbon face. Prepared grids were plunged into liquid ethane, clipped in an autogrid assembly and loaded in the electron microscope.

### X-ray Diffraction

Crystals were flash frozen in liquid nitrogen, with 10% ethylene glycol added as cryoprotectant. Diffraction patterns were collected in-house at 100 K with Cu Kα radiation (λ=1.5418 Å) using a CCD detector (Saturn 944++, Rigaku/MSC, Tokyo, Japan) mounted on a rotating anode generator (Micromax 007, Rigaku/MSC, Tokyo, Japan).

### Electron Diffraction Data Acquisition

#### Microscope Configuration and Adjustments

Electron diffraction experiments were conducted using a 200 keV Thermo Scientific Glacios® Cryogenic Transmission Electron Microscope equipped with a Ceta-D camera and a Falcon 4i Direct Electron Camera. Throughout our experiments, both the microscope stage and the autoloader were maintained at cryogenic temperatures (< 100 K). Microscope adjustments performed prior to data acquisition are described in detail in Supplementary Methods S2.

#### Diffraction Data Collection

Continuous rotation ED data were collected using the EPU-D software version 1.11, with a 50 μm C2 aperture to limit the beam size to 1.7 μm in the parallel beam nano-probe. Prior to acquiring diffraction patterns from a crystal, the eucentric height was adjusted at a nearby location, without exposing the crystal itself (see details in Supplementary Methods S2.2). At the same location, the direct [000] beam was finely centered in diffraction mode to hide it behind the beam-stopper. The beam was blanked, the stage moved mechanically to bring the crystal to the beam position (without changing any optical adjustments), and then the beam opened for immediate data recording.

For the elongated Gab1-SHP2 complex whiskers, multiple acquisitions were recorded along different sections of a single crystal (resulting in multiple diffraction datasets for each crystal), thereby minimizing the cumulative radiation damage to maintain a high signal-to-noise ratio. When possible, parts of the crystal located in the grid holes were preferred to avoid the background contribution of the grid film material. After collecting one segment of data, the stage was moved along the crystal, leaving at least 1 μm between the edges of the recording areas to avoid overlapping the exposure areas. Depending on the length, diffraction quality and the grid position of the crystal, up to fifteen data sets with 20° to 30° tilt segments (0.5°/frame) were acquired to cover the maximum possible tilt range of the sample stage (−70° to 70°) in overlapping segments (Table 1, Supplementary Table S3). A single acquisition with a higher tilt range of 60° (−30° to 30°) was recorded around the center of the tilt axis to determine the space group and unit cell parameters. High tilt angle segments (−70° to -50° and 50° to 70°, respectively) were collected multiple times, as several such segments could not be processed due to high background noise (resulting from the increased sample/ice thickness at high angles and/or beam path obstruction) and had to be excluded from the final dataset. The incident beam fluence during data collection was fixed and the data were collected with a total cumulative fluence of 2 to 4 e^−^/Å^2^ except for the acquisition with the higher tilt range collected with a total cumulative fluence of around 5 e^−^/Å^2^. Electron diffraction images were recorded as single frames in SMV format.

**Table 1:**
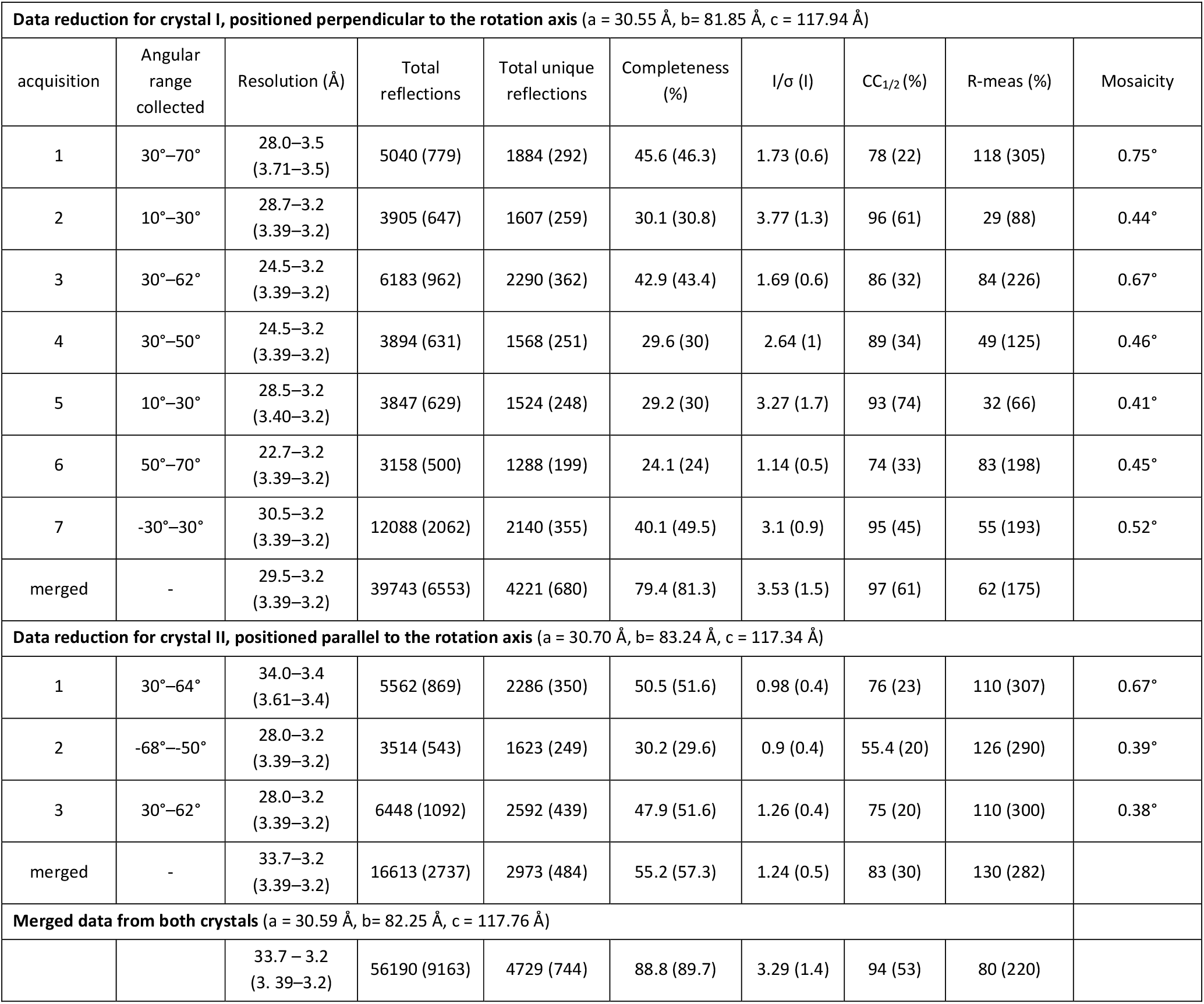
Data collection statistics for crystals used to determine the structure of the Gab1-SHP2 complex (space group P2_1_2_1_2_1_; all data collected at an accelerating voltage of 200 kV, corresponding to an electron wavelength of 0.025 Å).

#### Data Processing and Model Building

Data processing was carried out using the XDS software package [44]. The orthorhombic space group P2_1_2_1_2_1_ was identified from the central tilt range (−30° to 30°) with unit cell parameters a = 30.51 Å, b = 82.15 Å, c = 118.16 Å, and α = β = γ = 90°. These values were subsequently used to process datasets acquired at other tilt ranges. Datasets collected at multiple tilt ranges along the same crystal were merged in XSCALE [44], which were merged with corresponding data from isomorphous crystals (unit cell parameter differences of less than 1 %). Only individual tilt segments that contributed to improved quality (CC½), redundancy and/or completeness of the merged and scaled data were included in the final dataset. Diffraction data to a resolution of 3.2 Å with a completeness of 89 % could be obtained from just two crystals (Table 1), with averaged unit cell parameters a = 30.59 Å, b = 82.25 Å, c = 117.76 Å, and α = β = γ = 90° calculated in CELLPARM implemented in XSCALE. Merged data were exported in MTZ format using XDSCONV [44] and initial phases were determined by molecular replacement with PHASER [45] using the crystal structure of SHP2^1-222^ (PDB ID 5DF6 [46], Chain A) as search model (described in detail elsewhere [43]). Briefly, the search model was split into two ensembles, with ensemble 1 containing residues 4-104 (TFZ score 23.1) and ensemble 2 consisting of residues 108-220 (TFZ score 10.4). The molecular replacement solution was refined initially in REFMAC [47] in CCP4i2 software package [48] using scattering factors for electrons. The Coulomb potential map revealed density for the SHP2 linker between the two ensembles as well as residues of the bis-phosphorylated Gab1 peptide, which were fitted to the density using COOT [49] and refined with PHENIX [50] to 3.2 Å. The final model, consisting of SHP2 residues 4 to 221 and two Gab1 fragments (622-631 and 652-672), exhibits the following statistics: R_work_ / R_free_ = 29.9 % / 35.5 %, RMS bond lengths / angles 0.0045 Å / 0.58°, with 95.6 % / 4.4 % of the residues in the preferred / allowed regions of the Ramachandran plot. Figures were prepared using PyMol (The PyMOL Molecular Graphics System, ver. 2.0, Schrödinger, LLC).

## Results

The Gab1-SHP2 complex crystallized as very thin whisker-like crystals (Figure 1A) that did not diffract X-rays in-house, probably due to their small volume. Plate-like crystals that grew in the same drops (Figure 1B) did diffract X-rays to up to 2.7 Å resolution, but exhibited considerable disorder with multiple lattices (Figures 1C, D). 3D-ED/MicroED was therefore explored as a promising alternative for structure determination, given its potential to collect data from very small crystal volumes. In contrast to published protocols for lysozyme, several modifications had to be implemented during sample preparation, data acquisition, and data processing for the successful determination of the Gab1-SHP2 complex crystal structure. For vitrification, the single-sided blotting procedure with crystallization buffer applied to the back face (Supplementary Figure S3E) proved crucial to obtaining a sample containing visible crystals in thin ice.

**T**he cryo-EM system used here operates at 200 kV, which results in higher stopping power and inelastic scattering cross sections compared to 300 kV and therefore necessitates thinner specimens [51]. To assess the microscope performance, lysozyme was used as a well-studied sample (Supplementary Methods S1). Following the successful acquisition of diffraction data from tetragonal lysozyme crystals, the data collection procedure was applied to the Gab1-SHP2 sample. Gab1-SHP2 crystals were screened for diffraction behavior by recording single diffraction images without tilting the sample stage (tilt angle 0°) using a fluence of ∼ 0.1 e−/Å^2^. For crystals diffracting to a resolution of 3.5 Å or higher, datasets were collected within a total tilt range of 90° (−45° to 45°) using a total fluence of 8 e−/Å^2^ (Figure 2A). Rapid crystal deterioration was observed as the acquisition progressed, indicated by a rapid decrease in the resolution limit on consecutive frames (Figure 2B). Frames collected to a cumulative fluence of **⪅** 4 e−/Å^2^ revealed Bragg reflections to resolutions higher than 3.6 Å. Thereafter, the number of observed diffraction spots per frame decreased rapidly, with the resolution limit falling to ∼ 4.5 Å at a cumulative fluence of 6 e−/Å^2^ and to ∼ 6.5 Å at a cumulative fluence of 8 e−/Å^2^ (Figure 2B). It was not possible to index these data, precluding further processing.

**Figure 2:**
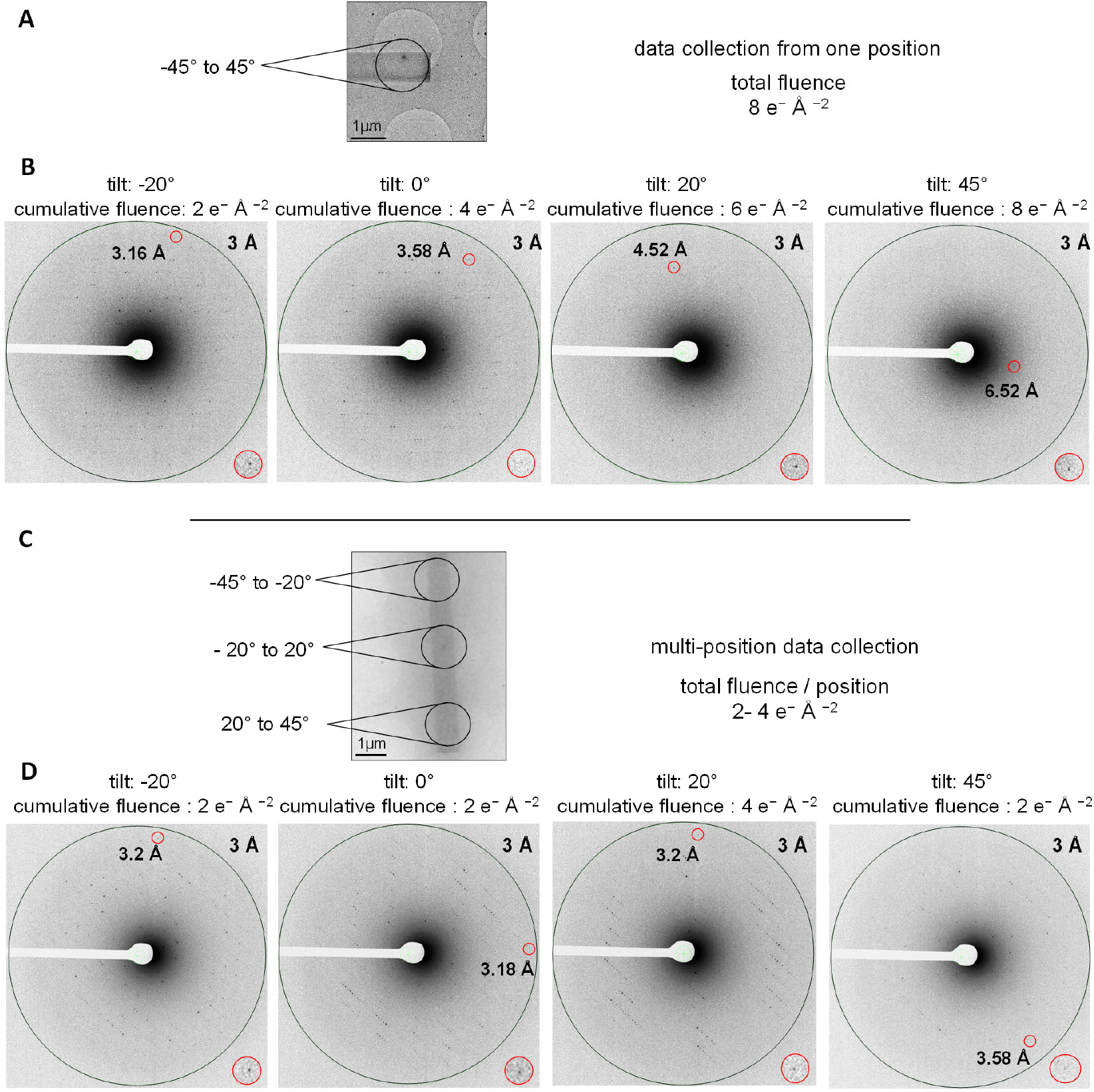
Multi-position data acquisition strategy reduces the cumulative incident beam fluence during 3D-ED/MicroED data collection. (A) Crystal used for 3D-ED/MicroED data collection at a single position while tilting the sample stage through a total range of 90°. (B) Resulting diffraction patterns at the indicated tilt angles with corresponding cumulative incident beam fluence. (C) Multi-position 3D-ED/MicroED data collection involves acquisition of diffraction data from multiple positions along a single crystal within an accumulated total tilt range of 90°. (D) Resulting diffraction patterns taken at the indicated tilt angles using a cumulative fluence ≤ 4 e−/Å^2^ throughout the acquisition range. For each diffraction image, the highest resolution Bragg reflection is highlighted with a red circle (magnified view lower right). Data recorded using the multi-position strategy show significantly improved diffraction, with distinct reflections up to around 3.2 Å resolution.

To minimize the cumulative incident beam fluence applied to the crystal during acquisition, two modifications were implemented to the data collection procedure. First, eucentric height adjustment prior to diffraction data collection was performed near the region of interest with a minimal fluence, avoiding exposing the crystal itself (detailed procedure in Supplementary Methods S2). Second, we took advantage of the elongated rod-like shape of the crystals. Instead of recording data from a single position, frames from consecutive tilt ranges were split over several positions along an individual crystal (Figure 2C), inspired by helical X-ray diffraction data acquisition strategies for needle-like crystals at synchrotron facilities [40], [41]. Specifically, the tilt range was divided into 3 intervals: -45° to -20, -20° to 20°, 20° to 45° (Figure 2C), with a wider tilt window around the zero-tilt angle to aid in indexing. Collecting fewer frames over smaller tilt ranges resulted in a reduction in beam exposure to the illuminated section of the crystal (and thereby radiation damage). Combining tilt series from different parts of the same crystal maximizes reciprocal space coverage while minimizing the effective total exposure over the entire acquired tilt range, resulting in data collected with a total cumulative fluence of 2 to 4 e−/Å^2^. This is a reduction in the total applied incident beam fluence by at least one half (Figure 2D), maintaining the diffraction quality of the crystal (∼ 3.2 Å) across the entire applied tilt range.

The data could be indexed in space group P2_1_2_1_2_1_ with unit cell parameters: a = 30.5 Å, b = 79.0 Å, c = 120.6 Å, and α = β = γ = 90°. Combining three segments collected from a single crystal over a total tilt range of 90° yielded data to a resolution of 3.2 Å with a completeness of 68% (Supplementary Table S3). Corresponding data from a second crystal were 61% complete, but turned out to represent the same region of reciprocal space (Supplementary Figure S4), indicating a similar orientation on the grid. In order to increase reciprocal space completeness systematically, data were collected from crystals positioned in orthogonal orientations on the grid, allowing the individual crystals to be tilted around different crystallographic axes. In addition, we utilized the maximum allowed sample stage tilt range (−70° to 70°). The latter can be performed only when the crystal has a clear path to the beam at high tilt angles that is not obstructed by grid bars, other crystals or contamination. Careful screening of the grids identified two adjacent crystals I and II that were positioned perpendicular to each other (Figure 3A). Due to the length of crystal I (oriented nearly perpendicular to the tilt axis), data collection could be performed several times in overlapping tilt intervals, increasing the quality and redundancy of the data and resulting in a completeness of 79 % at 3.2 Å resolution (Figure 3B, Table 1). High tilt angle data for crystal II oriented almost parallel to the tilt axis could be collected from three separate positions, resulting in a 55% complete data set to 3.2 Å resolution (Figure 3C). Merging the data from both crystals resulted in an overall completeness of 89 % at 3.2 Å resolution, with <I/σ(I)> / CC_1/2_ values of 3.29 / 94 % respectively (1.4 / 53 % in the highest resolution shell) (Table 1, Figure 3D). Coulomb potential maps calculated from the merged dataset are of improved quality, with better connectivity compared to the maps calculated from each crystal separately (Figure 3), displaying clear density for the Gab1 fragment bound to SHP2^1-222^ (Figures 3D, 4). The biological function and significance of the structure are presented elsewhere [43].

**Figure 3:**
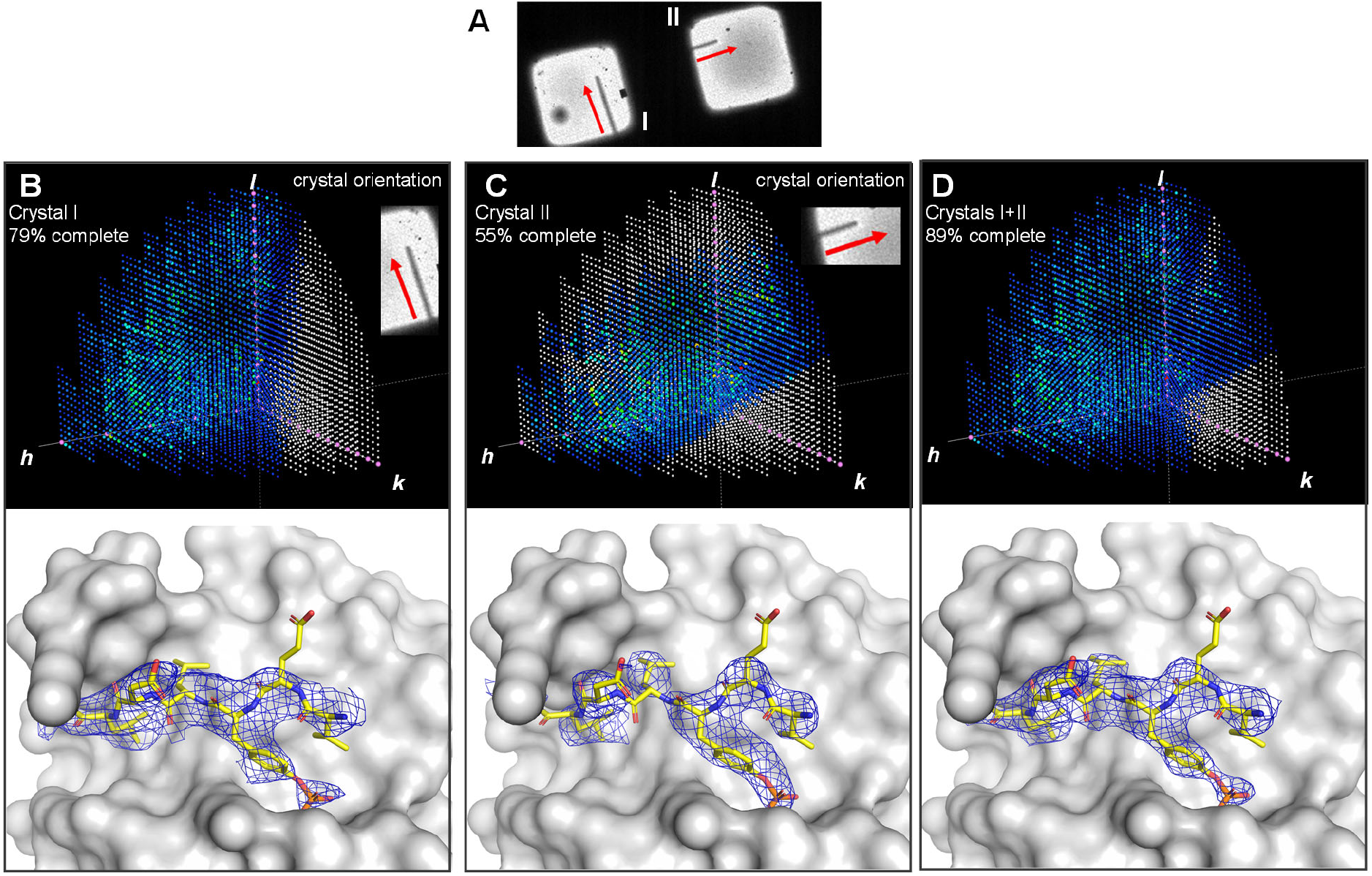
Improving data completeness by merging data collected from crystals of different orientations on the sample grid. (A) Zoomed in atlas view showing crystals I and II positioned perpendicular to each other on adjacent grid squares of the same sample. EM stage tilt axis vertical, red arrows show direction of consecutive data acquisition regions. (B-D) Effect of data completeness on the quality of calculated maps for the Gab1 fragment. Upper panels: Reciprocal space reflection distribution prepared with 3D Data Viewer in Phenix [50]. Missing reflections shown in white and systematic absences in pink; it is evident from the former that the ***b\**** axes of both crystals are in the direction of the electron beam. Lower panels: 2F_o_-F_c_ maps (contoured at 1.5 σ, blue mesh) for Gab1 residues 624-631 (yellow sticks) in complex with the N-terminal domain of SHP2^1-222^ (grey surface) using phases from the final structure refined against the corresponding data set. (B) Data collected from crystal I (oriented perpendicular to the rotation axis) with a total completeness of 79 % results in density for the peptide but misses a significant wedge of data. (C) Crystal II (oriented parallel to the rotation axis) yielded data with a completeness of 55 %, resulting in recognisable yet broken density for the peptide. (D) Merging the data from both crystals I and II reduces significantly the missing wedge, resulting in a better-defined density with improved connectivity.

**Figure 4:**
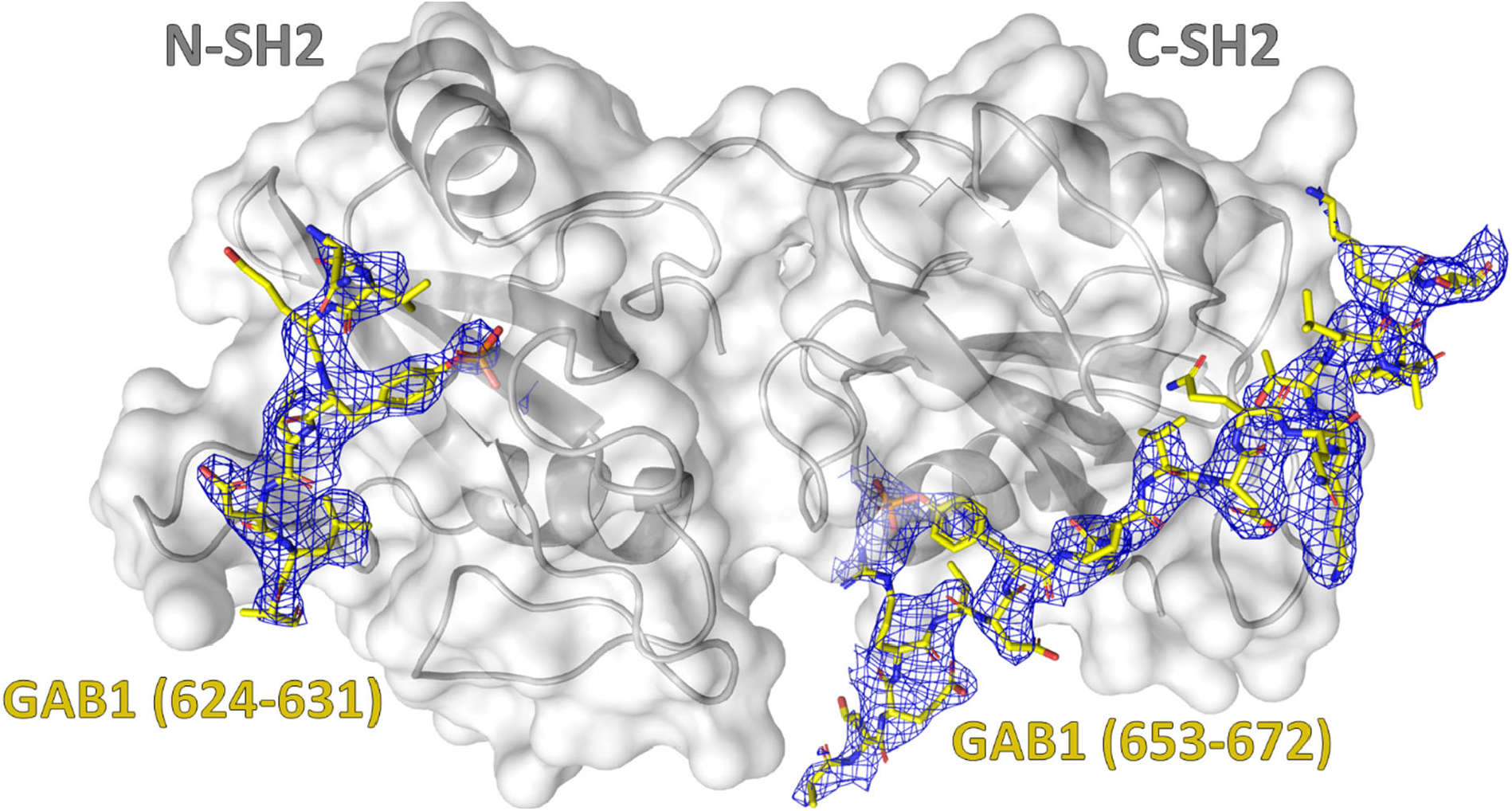
Crystal structure of the Gab1^617-684^ fragment in complex with SHP2^1-222^ [43] determined by 3D-ED/MicroED. SHP2 is depicted in cartoon representation (dark grey) with a light grey surface; ordered parts of the Gab1 peptide are shown as yellow sticks with corresponding 2F_o_-F_c_ density (contoured at 1.5 σ, blue mesh).

## Discussion

The procedures outlined in this paper were essential to determining the structure of the Gab1-SHP2 complex [43]. Although the crystals obtained were unsuitable for X-ray diffraction studies, either due to their thin needle-like form (yielding insufficient diffraction volume) or through crystal defects (splitting, twinning and/or high mosaicity) (Figure 1), they proved amenable to electron diffraction. Obtaining sub-micrometer-thick crystals embedded in a thin layer of vitrified ice [52] is essential to 3D-ED/MicroED as this allows electrons to penetrate the sample with minimal inelastic scattering [53]. Microcrystals can be grown by adjusting the crystallization conditions [54], [55] or obtained through fragmentation of larger crystals by e.g. crushing with a glass rod, vortexing with beads, or treating in a sonication bath [22], [25]. In the present case, microcrystals could be harvested directly from drops that contained macroscopic crystals without further preparation steps. Cryogenic focused ion beam (cryo-FIB) milling [56], [57] provides an alternative route to preparing crystals of ideal thickness, even for such challenging crystals as those obtained using lipidic cubic phases (LCP) [53], [58]–[61]; on the other hand, cryo-FIB-milling is currently not readily available in cryo-EM laboratories due to costly specialized instrumentation.

In the present study, it was necessary to adopt a single-sided back-blotting procedure prior to vitrification in order to obtain crystals on the EM grid. Back-blotting was also essential for vitrification of R2lox crystals [33], which similarly grew in a viscous (44% PEG 400) mother liquor. The simple modifications to the blotting setup introduced here (Supplementary Figure S3E) allow vitrification of crystals grown in high-viscosity buffers under controlled conditions of temperature and humidity. Another promising approach is the recently introduced pressure-assisted method, Preassis [27], which uses suction pressure to draw excess liquid through a sample grid resting on filter paper.

The crystals showed rapid deterioration even at relatively low beam fluence (Figure 2). It has been shown that electron-induced site-specific damage can occur at incident beam fluences as low as ∼0.9 e−/Å^2^ (disulfide bond cleavage) or ∼2.5 e−/Å^2^ (decarboxylation of acidic side chains) [19]. Application of the multi-position acquisition strategy reduced the cumulative incident beam fluence by half, enabling data collection as multiple tilt segments with continuous rotation. This provides more efficient data collection than e.g. SerialED [39] [22], yielding a higher coverage of reciprocal space per position and crystal as well as providing a sufficient wedge of data for indexing uncharacterized crystals. A similar segment-wise approach has been used to collect ED data from thin small molecule crystal needles [62], although small molecule crystals tend to be less susceptible to radiation damage due to tighter packing and an absence of bulk solvent. The data collection strategy implemented here is not limited to needle-like crystals, but can also be applied to other morphologies such as thin plates.

Through a systematic choice of crystals with recognizably different orientations, a data set with 89 % completeness could be obtained from just two Gab1-SHP2 crystals, compared to the thousands needed to obtain comparable completeness by SerialED. Nevertheless, some reflections remained inaccessible due to a preferred orientation of the crystals (each with the crystallographic ***b\**** axis perpendicular to the grid, Figure 3, Supplementary Figure S4). This could be alleviated through the use of a high-tilt angle sample holder capable of 360-degree rotation, developed recently for electron tomography [63], [64], [65].

As we were unable to index any of the X-ray diffraction patterns, it is not possible to determine whether the microcrystals used for ED data collection are the same form as the macroscopic needles or plates, only that the microcrystals diffract electrons. Hence, we are unable to say definitively whether the 3.2 Å ED dataset obtained here represents the ultimate resolution limit of these crystals. Nevertheless, the use of an energy filter [66] (which would decrease background noise due to removal of inelastically scattered electrons) together with a direct electron detector [67] (in contrast to the fiber optic coupled imaging sensor used here [68]) could increase the signal-to-noise ratio of the diffraction data and thereby the resolution.

Overall, the optimizations introduced in this study should find application in the study of a variety of challenging samples, including those with high viscosity or limited numbers of diffracting crystals. These improvements do not require specialized or costly instrumentation and can be implemented in standard cryo-EM laboratories. By addressing key issues in sample preparation, data collection, and processing, this work contributes to advancing the accessibility and application of ED for macromolecular structure determination.

## Acknowledgments

The authors would like to thank Dr. Thomas Monecke (Institute of Pharmaceutical Biotechnology, Universität Ulm) for valuable assistance in initial data processing. We are also thankful to Dr. Ioannis Skalidis (ZIK HALOmem, Martin Luther University Halle-Wittenberg) for suggesting the use of electron crystallography for structure determination and PD Dr. Annette Meister for advice on vitrification. This work was supported by the Federal Ministry for Education and Research (BMBF ZIK HALOmem, grant no. 03Z22HN23 to P.L.K. and grant no. 03Z22HI2 to M.T.S. and P.L.K. as well as CORONAmem, grant no. 03COV04 to M.T.S. and P.L.K.), the European Regional Development Funds (EFRE) for Saxony-Anhalt (grant no. ZS/2016/04/78115 to M.T.S. and P.L.K. and grant no. ZS/2024/05/187255 to P.L.K.), the German Research Foundation (DFG, project numbers 391498659, RTG 2467 and 514901783, CRC 1664), the European Union (Horizon Europe ERA Chair “hot4cryo”, project number 101086665 to P.L.K.), and the Martin Luther University Halle-Wittenberg.

## Author Contributions

AS, FH, and LM designed the experiment. LM conducted the complex preparation and crystallization. FLK, FH, and AS optimized the sample vitrification. FLK performed vitrification. FH carried out microscopic alignment, adjustments and implemented the diffraction protocols. FH and AS performed the data collection. AS, CP, and LM processed and analyzed the data. AS, CB and LM solved the structure and refined the model. AS and FH drafted the manuscript. SMF, PLK, and MTS supervised the project and revised the manuscript.

