## Supplementary Information for "Strategies for Mitigating Radiation Damage and Improving Data Completeness in 3D Electron Diffraction of Protein Crystals"

[1]Institut für Biochemie und Biotechnologie, [2]ZIK HALOmem, Biozentrum, [3]Institute of Molecular Medicine (Tumor Biology), [4]Charles-Tanford-Proteinzentrum, Martin-Luther-Universität Halle-Wittenberg, Halle (Saale), Germany; [5]Institute of Chemical Biology, National Hellenic Research Foundation, Athens 11635, Greece and [6]Mitteldeutsches Zentrum für Struktur und Dynamik der Proteine, Martin-Luther-Universität Halle-Wittenberg, Halle (Saale), Germany

[†]These authors contributed equally to this work

**Table of Contents**

Methods S1: Establishing workflow using lysozyme S2

S1.1: Lysozyme sample preparation S2

S1.2: Data collection and processing from lysozyme crystals S2

S1.3: Benchmarking the microscope performance using lysozyme S2

Table S1: Data collection and refinement statistics for lysozyme S3

Figure S1: Structure determination of tetragonal lysozyme crystals by 3D-ED/MicroED S4

Methods S2: Microscope adjustments S5

S2.1: General microscope alignment and calibration S5

Table S2: Data collection parameters in EPU-D S5

Figure S2: Flow chart of microscope adjustment and alignment before data acquisition S7

S2.2: Adjustments prior to data acquisition S8

Methods S3: Optimization of 3D-ED/MicroED Sample Preparation S9

Figure S3: Procedure for vitrification of Gab1-SHP2 complex crystals S10

Table S3: Data collection statistics for initial crystals of the Gab1-SHP2 complex S11

Figure S4: completeness of data collected from initial Gab1-SHP2 crystals S12

References S13

### Supplementary Methods S1: Establishing workflow using lysozyme

#### S1.1: Lysozyme sample preparation

Lyophilized lysozyme, purchased from Sigma-Aldrich (L6876; 90% purity), was dissolved in 20 mM sodium acetate pH 4.7 to a concentration of 80 mg/ml. Batch crystallization was prepared in a 1.5 ml tube by mixing 100 µl of enzyme solution with 100 µl of 80 mg/ml NaCl dissolved in 20 mM sodium acetate pH 4.7. Crystals were observed after incubating the mixture at 20°C for 2-3 h. For crystal fragmentation, the tube was placed in a sonicating water bath for 15 s at power setting 30% at 37 kHz (Elmasonic P30H, Singen, Germany). The suspension of microcrystals floating in the supernatant was used to make vitrified samples for electron diffraction. 3.5 µl of the crystal suspension was applied to Quantifoil ® R2/1 grids plasma treated in a PELCO easiGLOW glow discharging machine at 15 mA for 25 seconds under 0.4 mBar residual air pressure. Excess liquid was blotted for 12 s on both sides using a 595-grade ashless filter paper (ThermoFisher Scientific) as described in the main text.

#### S1.2: Data collection and processing from lysozyme crystals

Diffraction images were recorded within a total tilt range of 90° (-45° to 45°) per acquisition at 1°/frame. The electron incident beam fluence during the data collection was fixed to a nominal fluence of 0.088 $e^{-}/Å^{2}$/frame in the normal section and a total fluence of 8 $e^{-}/Å^{2}$ for the given rotation range. A dataset collected from a single crystal with a total tilt range of 90° was processed, indexed, and integrated using the XDS software package [1] to a resolution of 2.8 Å (Supplementary Table S1). Phases were obtained by molecular replacement in PHASER [2] using the tetragonal lysozyme structure (PDB ID 3J4G) as a search model [3]. The molecular replacement solution was refined in REFMAC [4] with iterative use of COOT [5] for model building.

#### S1.3: Benchmarking the microscope performance using lysozyme

Lysozyme microcrystals (Supplementary Figure S1) were screened for diffraction by bringing the crystal to the EH and recording a single diffraction at a tilt angle 0° and using a beam fluence of ~ 0.1 $e{-}/Å^{2}$. When a crystal diffracted to a resolution equal or better than 3 Å, a dataset was collected while rotating the sample stage continuously within a total tilt range of 90° (-45° to 45°) using a fluence of 0.088 e−/Å2 per frame and a total fluence of 8 $e{-}/Å^{2}$. The number of spots and the quality of the diffraction pattern varied during the course of the data acquisition, with the best diffractograms collected around the centre of the rotation axis (-10° to 10°) with highest resolution spots at ~ 2.4 Å (Supplementary Figure S1C). As data collection continued, the impact of accumulated radiation damage became evident through decay of the diffraction quality (reduced spot count and resolution), particularly in later frames. The effect of the applied fluence on the quality of the obtained diffraction observed here has been identified previously as a major challenge to electron crystallography [3], [6], [7]. Due to the high (tetragonal) symmetry and robustness of the lysozyme crystals investigated here, a major fraction of reciprocal space could be covered using this procedure. This resulting dataset could be processed to a resolution of 2.8 Å at 80.7 % completeness using frames collected from a single crystal (Supplementary Table S1, Figures S1D and S1E).

Supplementary Table S1: Data collection and refinement statistics for lysozyme; values in parentheses correspond to the highest-resolution shell.

| | |
|---|---|
| Accelerating voltage | 200 kV |
| Wavelength (Å) | 0.025 |
| Number of crystals | 1 |
| Total angular range collected (°) | 90 |
| **Data reduction** | |
| Space group | $P4_32_12$ |
| a, b, c (Å) | 78.6, 78.6, 37.8 |
| α = β = γ (°) | 90 |
| Resolution (Å) | 55.5 – 2.8 (3. 0–2.8) |
| Mosaicity (°) | 0.36 |
| Total reflections | 20009 (3858) |
| Total unique reflections | 2574 (458) |
| Completeness (%) | 80.7 (80.4) |
| I/σ (I) | 3.35 (1.64) |
| $CC_{1/2}$ (%) | 81 (65) |
| R-meas (%) | 137 (216) |
| **Model Refinement of merged acquisitions** | |
| $R_{work}/R_{free}$ (%) | 0.29/0.32 |
| RMSD bonds (Å) | 0.0077 |
| RMSD angles (°) | 1.32 |
| Ramachandran (%) | |
| (favoured, allowed, outlier) | 90.6, 9.4, 0.0 |

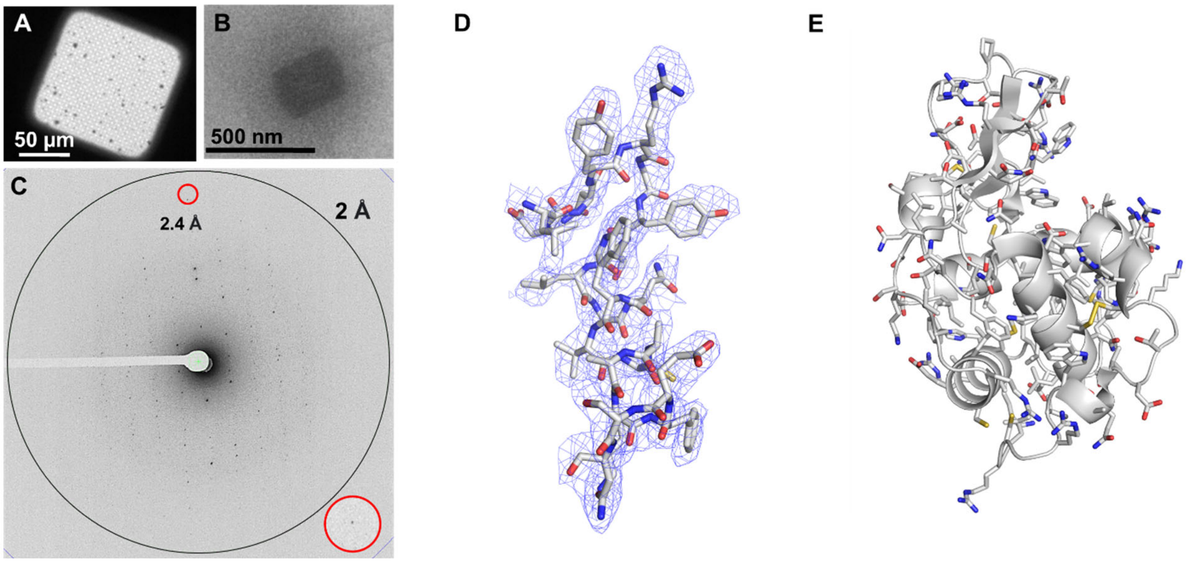


Supplementary Figure S1: Structure determination of tetragonal lysozyme crystals by 3D-ED/MicroED using a 200 keV Cryo-Transmission Electron Microscope equipped with a Ceta-D detector. (A) Low magnification cryo-EM view of lysozyme microcrystals on the sample grid square. (B) Zoomed-in view showing the crystal that was used for data set acquisition over a total tilt of 90°. (C) The diffraction pattern was recorded at a tilt angle 0° with a cumulative fluence of 4 e−/Å$^2$. (D) The resulting 2Fo-Fc Coulomb potential map around the residues 17-37 contoured at 1.5 σ together with (E) the complete final model.

### Supplementary Methods S2: Microscope adjustments

#### S2.1: General microscope alignment and calibration

To capture electron diffraction data effectively, we tuned the microscope to achieve the lowest possible incident beam fluence, preventing pixel saturation and blooming. To accomplish this, both the gun lens and C1 lens were set to their maximum values in our microscope (i.e., #8 and #11, respectively). The specific settings used with the EPU-D software are summarized in Supplementary Table S2.

Supplementary Table S2: Data collection parameters in EPU-D, all using Gun Lens number 8 and Ceta-D camera

| Preset | Optical Mode | Spot Number (C1) | Intensity (C2) | Magnification (Mag) Camera Length (CL) | C2 Aperture | Defocus |
|---|---|---|---|---|---|---|
| Diffraction Acquisition | Nano-probe (Diffraction) | 11 | Parallel Beam Value | 1.75 m (CL) | 50 μm | - |
| Imaging Acquisition | Nano-probe (SA) | 11 | Parallel Beam Value | 13500 X (Mag) | 50 μm | -5 μm |
| Search Auto-Eucentric | Microprobe (SA) | 1 | 0.600 20 μm Beam Diameter | 6700 X (Mag) | 50 μm | -30 μm |
| Gridsquare | Microprobe (LM) | 1 | 0.833 | 510 X | 50 μm | -100 μm |
| Atlas | Microprobe (LM) | 1 | 1.096 | 210 X | 150 μm | -500 μm |

To ensure minimal distortion in the diffraction pattern—particularly at higher resolutions—and to maintain a consistent, well-defined camera length across all datasets, we aligned the microscope for parallel beam illumination. The procedure for achieving a parallel beam is detailed in the following sections, especially the section (d).

Enhancing the signal-to-noise ratio (SNR) requires confining the diffracting electron beam within the crystalline domain. A larger beam introduces unwanted scattering from the amorphous carbon or vitrified water background, degrading the diffraction quality. There are two primary methods to achieve beam confinement: (a) Selected Area Electron Diffraction (SAED), which utilizes an aperture in the image plane of the objective lens to select a specific region of interest, and (b) nanoprobe mode, where the pre-field of the twin lens compresses the beam into a well-defined, smaller probe. We opted for nanoprobe mode over SAED for several reasons:

(i) SAED does not restrict the irradiated area, but rather selects electrons originating from a specific region of the sample. Consequently, the beam damage area extends beyond the diffracting domain, making it challenging to apply low-dose techniques.

(ii) SAED lacks precision, as the electrons contributing to the final diffraction pattern are not strictly confined to the selected region [8].

Another critical parameter is the beam diameter. While large beams introduce the amorphous background noise, excessively small beams reduce the diffracting domain, leading to a lower signal-to-noise ratio (SNR) and, in extreme cases, broadening of the diffraction spots. To balance these factors, we selected a beam size of 1.7 µm, corresponding to a 50 µm C2 aperture in nanoprobe mode on the microscope used in this study. Given the dimensions of our microcrystals, this beam size maximized the diffracting domain while minimizing background contributions from the amorphous substrate.

We followed the precise alignment and calibration procedure in the following sequence, using a sample of nanometric particles of thallium chloride on a lacey carbon support film grid (Agar Scientific S110):

(a) The microscope and sample were set up in the "Eucentric Focus" condition to ensure that the region of interest (ROI) on the sample stayed in focus and at the correct eucentric height (EH) within the imaging preset. To achieve this, precise EH adjustments were made using the standard stage tilt method within a ±30° range. Subsequently, the microscope was configured to the imaging preset, except for a temporarily lower spot number for better visibility. Following that, the focus of the objective lens was manually adjusted on the sample, which had previously been positioned at the EH, and the current value of the objective lens recorded as the EH-Focus setting.

(b) The EPU-D software was calibrated for the minimum image shifts at various magnifications using the standard method provided by the software [EPU-D User Manual Section 5.2].

(c) For lower magnification presets (*i.e.* Search/Auto-Eucentric, Gridsquare and Atlas), we performed beam and C2 aperture centering to prevent any image cutoff.

(d) In the nano-probe mode, specifically for Imaging and Diffraction Acquisition (with the twin-lens pre-field active), we initially aligned the microscope to achieve symmetrical parallel beam illumination around the optical axis [9]. The steps for such general microscope alignment are summarized in Supplementary Figure S2. As it is challenging to see the edge of the objective aperture at very low beam intensities, (i.e. at spot number 11 and gun lens 8), the edge of the aperture was brought into focus at a temporarily lower spot number with increased beam current. While objective fine alignments like astigmatism, current centering and coma-free alignment are not usually necessary for diffraction experiments, it is advisable to establish sound initial conditions, particularly for microscopes less commonly used for 3D-ED/MicroED.

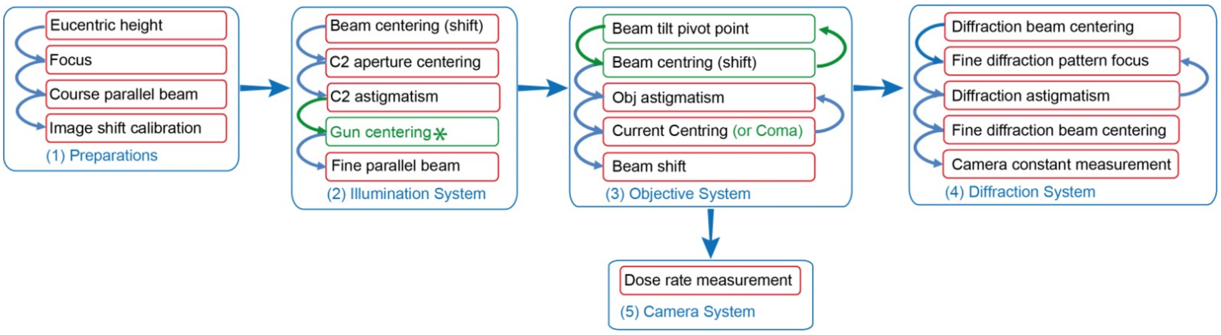


Supplementary Figure S2: Flow chart of microscope adjustment and alignment before data acquisition.

(e) We centered and fine-focused the diffraction pattern and adjusted the diffraction lens astigmatism. The microscope parameters in fully aligned and focused diffraction mode under nano-probe parallel beam conditions were then set as the diffraction acquisition preset in the EPU-D software. The values of the astigmators, deflectors, and optical state were recorded separately as an alignment file and gun register. Particular attention was paid to recording the values of diffraction and objective lenses, ensuring precise diffraction and image focus.

(f) We calibrated the camera constant, and subsequently, the camera length based on the 200 kV accelerating voltage, using the same thallium chloride sample. For this purpose, we recorded several diffraction patterns and measured the positions of numerous Friedel pairs to determine the beam centre and respective 2θ angles. In this study, the camera length on the Ceta-D camera, at the nominal camera length of 1750 mm, was calculated to be 1704.5 mm, equivalent to a camera constant of $3.0544 \times 10^3$ pixel.Å. Although it is possible to determine the camera length based on the final data, a high-fluence diffraction pattern from a hard material yields a more accurate measurement.

We measured the fluence at each preset meticulously to be able to track the dosage applied to each crystal. Accurate quantification of the fluence can be challenging, particularly when using the Ceta-D camera in diffraction mode or when the aperture or grid bars partially obstruct the beam. As our microscope lacks a Faraday cup, we employed the Falcon III EC direct electron detector in counting mode for precise fluence quantification. Ensuring that the microscope was in the previously aligned state, the sample was removed and the magnification adjusted manually, leaving all other parameters unchanged. Once the fluence fell within the "green zone" (indicating an acceptable range) in the Falcon III EC reference image manager software associated with the direct electron camera, a 10-second exposure image in counting mode was captured and the average fluence from the resulting image was calculated.

Immediately prior to each data collection session, the camera gain reference was updated using the standard protocol with the CetaD reference image manager software to ensure the accuracy of our data acquisition process. The microscope was now ready for loading vitrified grids containing submicrometer protein crystals.

**S2.2: Adjustments prior to data acquisition**

Before acquiring the diffraction patterns of each crystal, the EH was re-adjusted. Since the optimal fluence limit of the electron diffraction data is very small [10], we decided to adjust the EH with a minimal possible incident beam fluence by adjusting the height at a nearby location rather than on the region of interest (ROI) of the crystal. One of the following methods was used for such "Low fluence EH adjustments":

(a) When crystals were evenly distributed and well-separated on the grid and the grid square appeared flat and undamaged, a suitable empty spot for adjusting the EH was easily located using standard procedures. In such instances, the stage was moved to navigate to a vacant area nearby (ca. 15 μm) and the EH fine-tuned in the EPU-D software's search/ Auto-Eucentric preset using the Auto-Eucentric Height function.

(b) In cases where approach (a) was not feasible, e.g. due to numerous crystals, contaminations near the ROI, or difficulties arising from thick ice, the EH was adapted at a higher magnification and a smaller beam diameter, essentially the imaging preset. To achieve this, the objective lens was set to the EH-Focus value as explained earlier, and the stage Z movement was used to focus the region ≤2 μm from the edge of the crystal.

In both scenarios, the EH adjustment was checked by capturing very low-fluence and low-magnification images of the targeted crystal region with a ±30° tilt (or at the maximum tilt range if it was of a larger value) to ensure that the crystal remained in the field of view and nothing obstructed the beam during the tilt.

Following these adjustments, the primary beam position was determined on the fluorescent screen at the grid position used to make the EH adjustment (empty carbon film). This allowed precise placement of the beam-stopper to conceal the central beam, preventing camera saturation and pixel blooming while preserving the clarity of fine diffracting spots. At this point, the microscope was ready to capture high-quality diffraction patterns. The beam was blanked and the stage was adjusted mechanically to position the ROI of the crystal under the beam, with no changes to the optical adjustments. After unblanking the beam, data recording was started immediately to minimize any potential beam drift.

### Supplementary Methods S3: Optimization of 3D-ED/MicroED Sample Preparation

Using the same procedures for the Gab1-SHP2 sample as for lysozyme resulted in grids covered by a thick layer of ice, obscuring most of the grid squares (Figure S3B). The few grid squares visible within the blotting area exhibited rounded edges, indicating relatively thick ice even in the visible area. In contrast to tetragonal lysozyme, which crystallizes in a salt-based low-viscosity precipitant solution, the Gab1-SHP2 complex crystallized in 29% PEG3350, rendering the sample substantially more viscous. Increasing the blotting time to 24 seconds resulted in a non-uniform gradient of vitrified ice thickness throughout the sample, with more visible grid squares (Figure S3C). Although grid holes could be seen, indicating areas of thinner ice layer, no protein crystals could be identified in the blotted sample. To test whether microcrystals were damaged during sonication, crystal fragmentation was tried using a glass rod and vortexing with seed beads. Nevertheless, no crystals were identified in any of the prepared samples. To avoid the possibility that crystals were lost during the fragmentation process, crystal suspensions were pooled from the hanging drops and applied directly to the grid without any further treatment. Once again, no crystals were observed in any of the grid squares.

To investigate whether microcrystals may have been lost due to adhesion to the filter paper, back face blotting was explored [11], [12]. As the Vitrobot does not provide this possibility directly, the front face filter paper was replaced with an unstretched piece of parafilm that was cut to match the size and shape of the filter paper (Figure S3E). Initial attempts at back face blotting resulted in an opaque sample, indicating thick ice. A 3.5 µl drop of crystallization buffer diluted 1:1 with water was applied to the back face of the grid, reasoning that this could establish a connection between the viscous mother liquor of the crystals and the filter paper to allow efficient withdrawal of excess liquid through the sample grid holes. Implementation of both simple yet efficient modifications resulted in a gradient of thin ice across the sample grid (Figure S3D). Most grid squares were clearly visible with sharp and faceted edges, especially at the center of the grid and in the direction of thinner ice. In particular, thin crystals ranging in width from 0.5 µm to a few micrometers of diverse lengths were observed across the sample grid at various magnifications, some extending along several grid holes and even across several grid squares (Figures S3D, F). The grid was carefully screened to select crystals with a thickness suitable for electron diffraction data collection, several of which proved promising in diffraction mode (Figures 3A, C). Several grids of the Gab1-SHP2 complex crystals clipped in an autogrid assembly and loaded in the electron microscope. The proper ice thickness and presence of crystals were achieved only when both introduced modifications were applied during sample preparation. It should be noted that crystal thicknesses were not measured directly in this work, but rather indirectly based on transparent contrast, faceted edges, and quality of diffraction at zero-degree tilt.

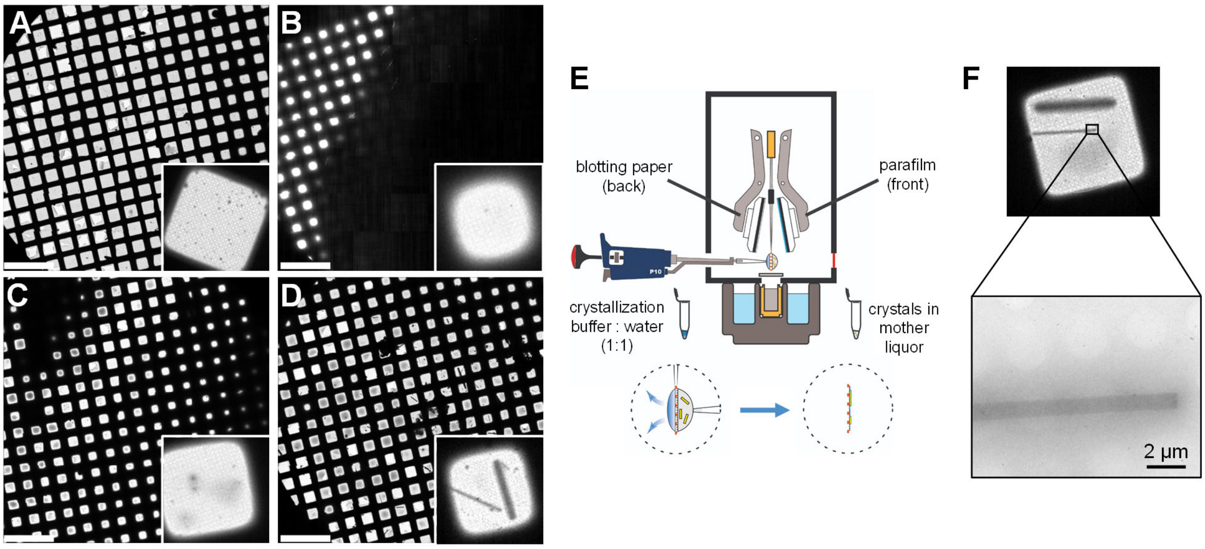


Supplementary Figure S3: Procedure for vitrification of Gab1-SHP2 complex crystals using the Thermofisher Vitrobot Mark IV. (A-D) All-grid atlas views of crystal specimens (scale bars 450 µm) with zoomed-in views of corresponding single grid squares (insets, box edges 150 µm). (A) Tetragonal lysozyme crystal sample (non-viscous slurry) prepared by double-sided blotting for 12 s at 4 °C and 95 % humidity. (B) Gab1-SHP2 crystal sample (viscous slurry containing 29% PEG3350) prepared under conditions equivalent to (A). (C) Gab1-SHP2 sample upon increasing the double-sided blotting time to 24 s. (D) Gab1-SHP2 specimen prepared using the optimized setup (see (E)). (E) Schematic drawing demonstrating the optimized vitrification setup for preparing protein crystals for electron diffraction experiments. Complex crystals were harvested from the crystallization drops and applied directly to the front face of the EM grid. Crystallization buffer, diluted 1:1 with water, was applied to the back face of the grid. The plunger was mounted with a parafilm disc on the side facing the crystals, and with blotting paper on the side facing the buffer solution. (F) Low magnification cryo-EM image of Gab1-SHP2 complex crystals observed after sample vitrification using the optimized setup. Two long rod-shaped crystals are visible on the grid.

Supplementary Table S3: Data collection statistics for initial crystals of the Gab1-SHP2 complex (space group $P2_12_12_1$, α = β = γ =90°; all data collected at an accelerating voltage of 200 kV, corresponding to an electron wavelength of 0.025 Å).

| **Data reduction for crystal A** (a = 30.5 Å, b= 79.0 Å, c = 120.6 Å) | | | | | | | | | |
|---|---|---|---|---|---|---|---|---|---|
| acquisition | Angular range collected | Resolution (Å) | Total reflections | Total unique reflections | Completeness (%) | I/σ (I) | $CC_{1/2}$ (%) | R-meas (%) | Mosaicity |
| 1 | -45°–-20° | 16–3.2 (3.50–3.2) | 4766 (1149) | 1554 (361) | 29.5 (30.1) | 3.49 (1.11) | 96 (44) | 35 (133) | 0.38° |
| 2 | -20°–20° | 16–3.2 (3.50–3.2) | 7845 (1906) | 1645 (373) | 31.2 (31.1) | 1.31 (0.8) | 40 (22) | 83 (150) | 0.44° |
| 3 | 20°–45° | 16–3.2 (3.50–3.2) | 4654 (1116) | 2005 (474) | 38.1 (40) | 2.28 (0.59) | 92(31) | 48 (202) | 0.55° |
| merged | - | 17–3.2 (3.50–3.2) | 17256 (4173) | 3558 (815) | 67.7 (68) | 2.41 (0.92) | 53 (31) | 75 (161) | |
| **Data reduction for crystal B** (a = 30.60 Å, b= 80.07 Å, c = 120346 Å) | | | | | | | | | |
| 1 | -45°–-20° | 17–3.2 (3.50–3.2) | 4657 (1124) | 1734 (404) | 33 (33.7) | 2.3 (0.53) | 96 (9) | 48 (275) | 0.52° |
| 2 | -20°–20° | 17–3.2 (3.50–3.2) | 7977 (1906) | 1695 (384) | 32 (32) | 1.3 (0.94) | 23 (10) | 91 (116) | 0.45° |
| 3 | 20°–45° | 17–3.2 (3.50–3.2) | 4850 (1163) | 1797 (418) | 34 (35) | 1.13 (0.84) | 21 (20) | 48 (202) | 0.31° |
| merged | - | 17–3.2 (3.50–3.2) | 17492 (4190) | 3213 (734) | 61 (61.3) | 1.5 (0.86) | 32 (14) | 93 (134) | |
| **Merged data from both crystals** (a = 30.55 Å, b= 79.50 Å, c = 120.48 Å) | | | | | | | | | |
| | | 17 – 3.2 (3.35 –3.2) | 34748 (8363) | 3631 (831) | 69 (69.4) | 2.57 (1.12) | 58 (24) | 92 (147) | |

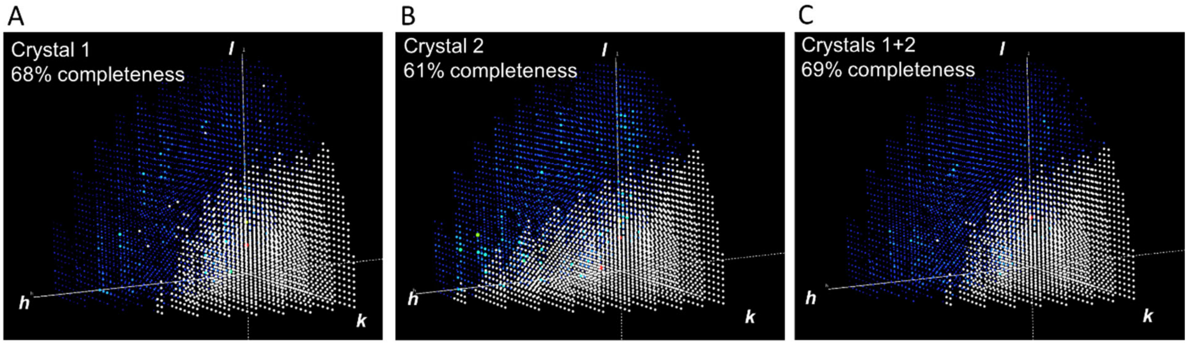


Supplementary Figure S4: Completeness of data collected from initial GAB1-SHP2 crystals. 3D reciprocal space representation of data collected from the first crystal in (A). Corresponding data from a second crystal (B) were 61% complete but represented the same region of reciprocal space, indicating a similar orientation on the grid; the merged data from both crystals showed no increase in reciprocal space coverage (C). Missing reflections are depicted in white. 3D representation of reflection spheres was prepared using 3D Data Viewer in Phenix [13].
